# Incorporating models of subcortical processing improves the ability to predict EEG responses to natural speech

**DOI:** 10.1101/2023.01.02.522438

**Authors:** Elsa Lindboom, Aaron Nidiffer, Laurel H. Carney, Edmund Lalor

## Abstract

The goal of describing how the human brain responds to complex acoustic stimuli has driven auditory neuroscience research for decades. Often, a systems-based approach has been taken, in which neurophysiological responses are modeled based on features of the presented stimulus. This includes a wealth of work modeling electroencephalogram (EEG) responses to complex acoustic stimuli such as speech. Examples of the acoustic features used in such modeling include the amplitude envelope and spectrogram of speech. These models implicitly assume a direct mapping from stimulus representation to cortical activity. However, in reality, the representation of sound is transformed as it passes through early stages of the auditory pathway, such that inputs to the cortex are fundamentally different from the raw audio signal that was presented. Thus, it could be valuable to account for the transformations taking place in lower-order auditory areas, such as the auditory nerve, cochlear nucleus, and inferior colliculus (IC) when predicting cortical responses to complex sounds. Specifically, because IC responses are more similar to cortical inputs than acoustic features derived directly from the audio signal, we hypothesized that linear mappings (temporal response functions; TRFs) fit to the outputs of an IC model would better predict EEG responses to speech stimuli. To this end, we modeled responses to the acoustic stimuli as they passed through the auditory nerve, cochlear nucleus, and inferior colliculus before fitting a TRF to the output of the modeled IC responses. Results showed that using model-IC responses in traditional systems analyses resulted in better predictions of EEG activity than using the envelope or spectrogram of a speech stimulus. Further, it was revealed that model-IC derived TRFs predict different aspects of the EEG than acoustic-feature TRFs, and combining both types of TRF models provides a more accurate prediction of the EEG response.x

## Introduction

Decades of research have sought to understand how the human brain processes the many sounds we encounter in everyday life. For example, since the 1930s researchers have used the electroencephalogram (EEG) to derive event-related potentials (ERPs) by averaging responses immediately following repeated presentations of brief, isolated stimuli (Davis, 1939; Handy 2005, Sur and Sinha, 2009). However, most sensory information available to human listeners is continuous, non-repeated, and occurs within a noisy environment, forcing listeners to discern important on-going signals from surrounding irrelevant information with only a single presentation. This includes that all-important of human signals – speech. To better approximate normal listening conditions – with a particular emphasis on speech – research turned towards using longer, more natural stimuli to elicit the EEG (Connolly et al., 1994; Näätänen, 1997). Initially, these studies used relatively short segments of speech to produce ERPs, which still provided only a limited view on how the brain parses and processes continuous segments of acoustically, lexically, and semantically rich speech.

In recent years, researchers have increasingly emphasized the use of continuous, natural speech in their experiments (Hamilton and Huth, 2020). One fruitful approach to analyzing the resulting neural data involves modeling those data based on the speech stimuli that elicited them (Brodbeck and Simon, 2020). This approach, which is known as system identification, treats the brain as something of a ‘black box’ and seeks to develop quantitative mappings between various speech features and the resulting neurophysiological responses. In particular, electroencephalography (EEG) has often been the recording modality of choice given its noninvasive nature, ease of use, and high temporal resolution (Gevins et al., 1995; Regan, 1989; Murakami and Okada, 2006; Buzsaki et al., 2012; Lopes da Silva and Niedermeyer, 2005). One particularly popular and tractable analysis involves treating the brain as a linear time-invariant (LTI) system and obtaining a so-called temporal response function (TRF) via regularized linear regression (Crosse et al., 2016, 2021). This framework allows researchers to study how the brain processes speech at different hierarchical levels by modeling the relationship between EEG and both acoustic (e.g., acoustic envelope, spectrogram) and linguistic (e.g., phonetic features, semantic surprisal) features (Di Liberto et al., 2015, Broderick et al., 2018; Brodbeck et al., 2018, 2022; Gillis et al., 2021). For example, a model involving spectrogram and phonetic features out-performs either constituent model, indicating that each feature contributes to unique aspects of the EEG signal (Di Liberto et al., 2015).

One limitation of the TRF approach – as noted by Drennan and Lalor (2019) – is that it makes a strong assumption about linearity and time invariance. In effect, this assumes that EEG responses to a particular speech feature always have the same timing and morphology; because the speech feature changes in intensity, the EEG response will scale, but will not change in terms of its timing or shape. However, this assumption is incorrect; it has long been known that EEG responses to auditory stimuli vary in both amplitude and latency with the intensity of the sound (Beagley and Knight, 1967). Drennan and Lalor (2019) proposed to relax this assumption in the context of modeling EEG responses based on the speech envelope. Specifically, they allowed the TRF to vary in morphology for different envelope intensities by binning the speech envelope based in amplitude deriving a multivariate TRF (mTRF). The resulting amplitude-binned (AB) envelope mTRF produced significant improvements in the ability to predict EEG responses to novel stimuli (Drennan and Lalor, 2019).

While this approach produced significant improvements, it was based on a simple, somewhat arbitrary manipulation of the stimulus (amplitude binning). A more principled approach not yet considered would be to formally incorporate the substantial processing of sound input that occurs along subcortical pathways before reaching cortex and contributing to scalp-recorded EEG. For example, it is well known that neurons in the inferior colliculus (IC) are tuned to sound frequency and amplitude-modulation rate (Krishna and Semple, 2000; Nelson and Carney, 2007). Thus, IC neurons represent a population of cells that can integrate responses to sound features (e.g., spectrogram, amplitude envelope) extracted at lower levels of the auditory system (e.g., Carney et al., 2015). Given that such a transformed representation of the speech input is what cortex actually receives (rather than the stimulus itself), incorporating a model of such subcortical processing might lead to improved predictions of cortical EEG.

In the present study, we hypothesized that accounting for subcortical processing of speech sounds would improve predictions of EEG responses to natural speech. To test this, we modeled IC responses to speech sounds using the phenomenological same-frequency, inhibitory-excitatory (SFIE) model based on Nelson and Carney (2004; Fig. 1). This model transforms a sound input into simulated responses at the levels of the auditory nerve (AN), ventral cochlear nucleus, and inferior colliculus that have been validated against neurophysiological recordings across a series of studies (Zilany et al., 2009, 2014; Nelson and Carney, 2004; Carney et al., 2015; Carney and McDonough, 2019). We fitted TRF models to these simulated IC responses, broadband speech envelopes, AB envelopes, and spectrograms and measured how well each TRF could predict EEG responses recorded while participants listened to speech. Overall, the mTRF fit to IC responses produced more robust EEG prediction than the speech envelope or spectrogram. The IC response and AB-envelope mTRFs performed at comparable levels. However, analysis of the correlations between predicted EEGs from these two models revealed that they predicted different aspects of the EEG. Thus, combining the IC-response and AB-envelope mTRFs further improved the EEG predictions.

**Figure 1:**
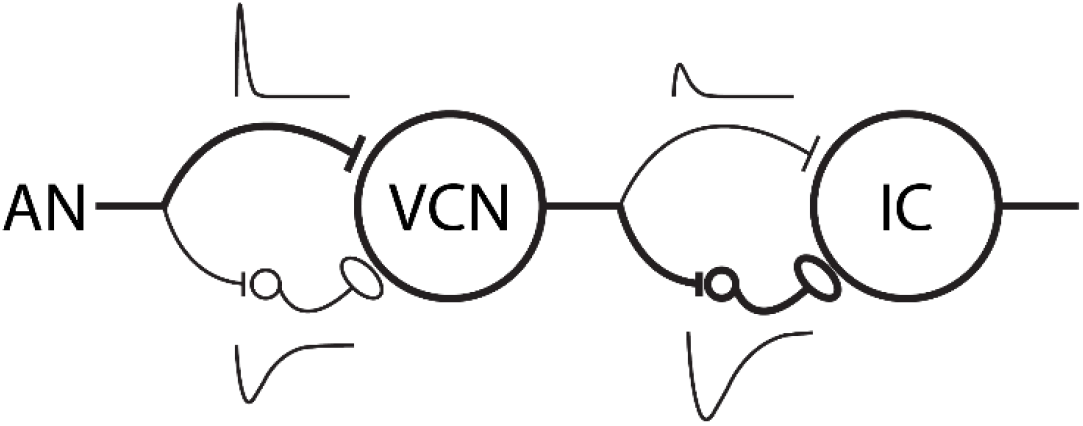
Schematic diagram of the same-frequency inhibition and excitation (SFIE) model. A single model AN fiber provides the postsynaptic cell with both excitatory and inhibitory input, via an inhibitory interneuron. The thickness of the lines corresponds to the relative strength of the inhibition and excitation at each level. Alpha functions representing the assumed membrane and synaptic properties are also shown above or below corresponding synapses. Adapted from Nelson and Carney (2004).

## Methods

### EEG Data and Stimuli

Stimuli and corresponding EEG responses from 19 subjects were obtained from two previous studies (DiLiberto et al., 2015; Broderick et al., 2018). In those experiments, subjects were presented with 20 three-minute-long segments of audio from an audiobook read by a male American English speaker (*The Old Man and the Sea* by Ernest Hemingway), using Sennheiser HD650 headphones. Each segment was ∼155 s in duration and segments were presented in sequential order. During stimulus presentation, 128 scalp channels (+ 2 mastoid channels) of EEG data were recorded from each participant with a sampling rate of 512 Hz using the BioSemi ActiveTwo system. The recordings were digitally filtered between 1 and 15 Hz with a Chebyshev Type 2 filter and referenced to the average of the two mastoid channels (as described in DiLiberto et al., 2015). After collection, the EEG signal was down sampled to 128 Hz to decrease computation time during further analyses.

### Speech Representations

We derived TRFs based on four distinct representations of the speech stimulus. These representations were presented as either single or multivariate feature vectors. Before feature extraction, all audio samples were lowpass filtered with a Chebyshev Type 2 filter having a cutoff frequency of 4.8 kHz. The broadband-envelope representation of the audio was calculated as

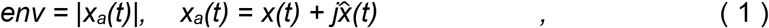

where *x*_*a*_*(t)* is the analytical representation of the signal, taken as the sum of the original speech, *x(t)*, and its Hilbert transform 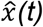 (Fig. 2A).

**Figure 2:**
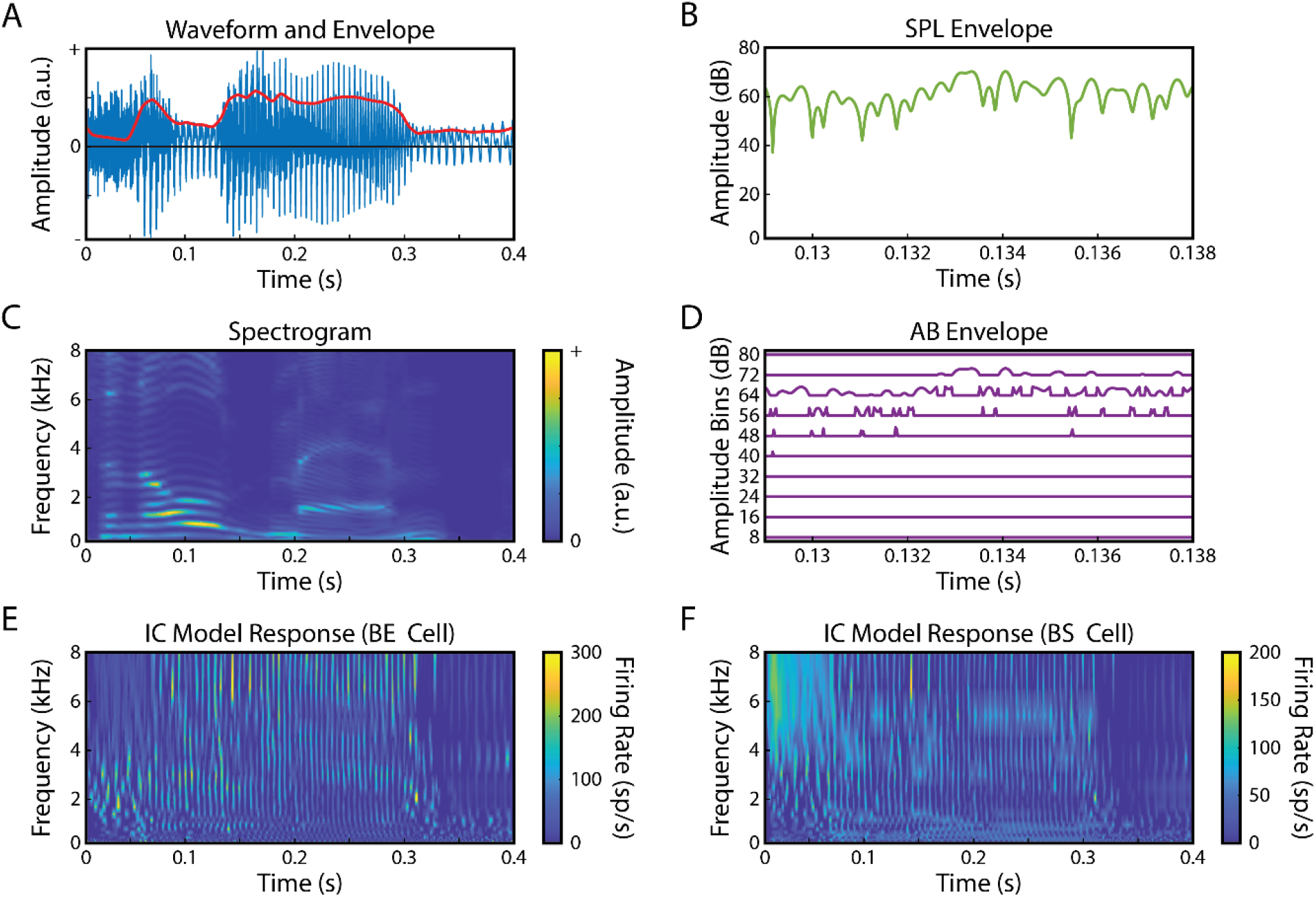
Representations of speech stimulus. Four different representations of the speech stimuli were used to derived TRFs and predict EEG responses: **A)** broadband envelope, **B)** SPL envelope used for amplitude binning, **C)** spectrogram, **D)** amplitude-binned envelope and **E, F)** IC BE and BS model responses.

The spectrogram of the speech was obtained by filtering the speech into 20 log-spaced frequency bands ranging from 200 to 8-kHz (Di Liberto et al., 2015). The amplitude envelope of each frequency band was calculated using Eq. 1 (Fig. 2C). All audio-feature signals were down sampled to 128 Hz to match the sampling rate of the EEG data. To create the AB-envelope feature vector, the SPL (sound pressure level) envelope (Fig. 2B) was binned into 10 8-dB level ranges using the histcounts function in MATLAB (as outlined in Drennan and Lalor, 2019).

This binning resulted in 10-variable feature vectors at each time point (Fig. 2D). Neural responses to the acoustic stimuli were modeled in two steps. First, an AN model (Zilany et al., 2014) was used to simulate the responses of 20 AN fibers with characteristic frequencies (CF, the frequency that elicits a response at the lowest sound pressure level, SPL) that were matched to the spectrogram frequency bands. Then, the SFIE model (Carney and McDonough, 2019) was used to simulate responses of two types of IC neurons: band-enhanced (BE) and band-suppressed (BS) neurons, as described by modulation transfer functions, average rates as a function of modulation frequency in response to sinusoidally amplitude-modulated sounds (Kim et al., 2020). BE IC neurons are excited by amplitude-modulated stimuli with modulation frequencies near the peak of the modulation transfer function (MTF), whereas BS IC neurons are suppressed by stimuli that are modulated near a trough frequency in the MTF. The IC models had peak or trough modulation frequencies in the MTFs that were set to 100 Hz, which is near the center of the distribution for MTFs recorded in the IC (Kim et al., 2020). The IC-model feature vector consisted of responses from 20 BE and 20 BS neurons, with CFs ranging from 200 Hz to 8 kHz to match the frequencies used for the spectrogram analysis (Fig. 2E, F). BE and BS responses were concatenated, resulting in a 40-variable feature vector at each time point. The responses of the neural models were also down sampled to 128-Hz for correlation analysis with the EEG signals. Code for the AN and IC models is available at https://urhear.urmc.rochester.edu.

### TRF calculation and EEG prediction

The TRF is a linear transformation from a stimulus feature vector, *S(t)*, to the neural response vector, *R(t)*, i.e.,

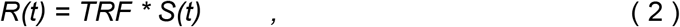

where * represents the convolution operator (Crosse et al., 2016). The TRF for each feature is calculated over a series of time lags between the stimulus and the response, producing a set of temporal TRF weights for each EEG channel. To estimate the TRF we used ridge regression (see Crosse et al., 2016 for details on the TRF). In brief, this involves solving for the TRF using the following equation:

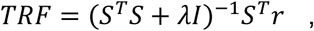

where *S* is the lagged time series of the stimulus property, *s*(*t*), and is defined as follows:

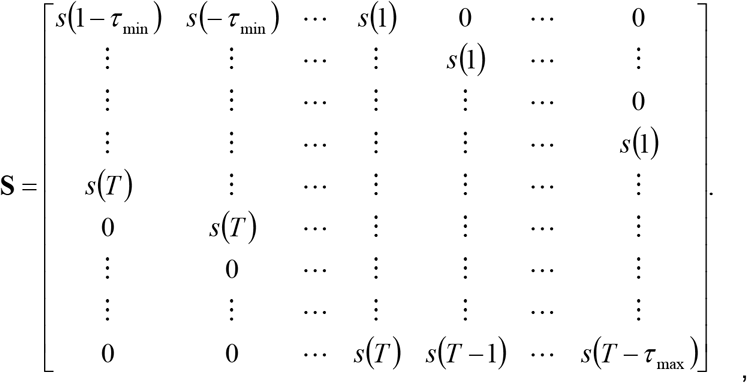

where the values *τ*_min_ and *τ*_max_ represent the minimum and maximum time lags (in samples), respectively. In S, each time lag is arranged column-wise and non-zero lags are padded with zeros to ensure causality (Mesgarani, et al., 2009). The window over which the TRF is calculated is defined as *τ*_window_ = *τ*_max_ – *τ*_min_ and the dimensions of *S* are thus *T* × *τ*_window_ (where *T* is the total length of the stimulus/data used for fitting). To include the constant term (y-intercept) in the regression model, a column of ones is concatenated to the left of *S*. The neural response data is organized into a matrix *r* with the *N* EEG channels arranged column-wise (i.e., a *T* × *N* matrix). The resulting TRF, *w*, is a *τ*_window_ × *N* matrix with each column representing the univariate mapping from *s* to the neural response at each channel. *λ* is a regularization parameter that controls for overfitting (see below for how this was determined). TRFs were estimated separately for each stimulus feature representation. Initially, time lags were set to - 500 to 500 ms, but the window of analysis was narrowed to 0 to 275 ms after inspection of TRFs (Fig. 3).

**Figure 3:**
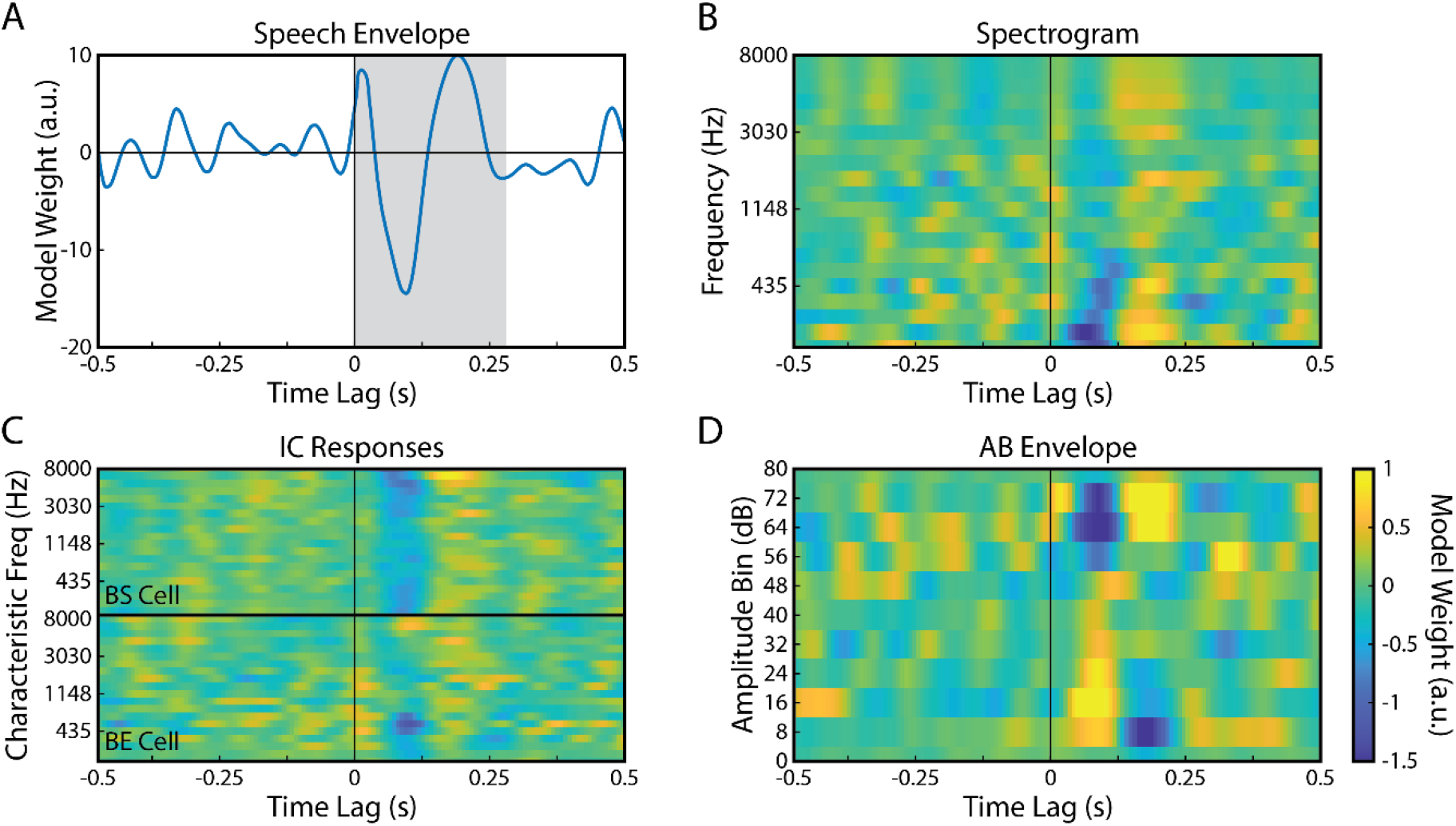
TRF model weights plotted over wide range of time-lags. mTRF plotted for the **A)** Envelope, **B)** Spectrogram, **C)** IC responses provided by SFIE model; top half shows IC BS responses; bottom half shows IC BE responses, and **D)** AB envelope representations of the speech stimulus. Time lags ranged from -500 to 500-ms and mTRFs were averaged across the 12 electrodes of interest (see Fig. 4). Shaded region in **A)** indicates the 275-ms analysis window used.

A cross-validation step was completed to assess the EEG’s sensitivity to different speech features as well as to select the optimal ridge parameter to prevent overfitting (a detailed description of this process is provided in Crosse et al., 2016). To summarize, ‘leave-one out’ cross-validation was used in which TRF models were fit to a group of trials for each ridge-parameter value (λ = 10^−6^,10^−4^, 10^−2^…10^6^). TRFs from all but one trial were averaged and used to predict responses from the remaining trial. This process was repeated until all trials had been left out and until all ridge parameters within the predetermined range were exhausted. The best ridge parameter was chosen based on the correlation value between predicted and measured EEG data. This regularization parameter was found using ‘mTRFcrossval.m’ and was optimized for each stimulus and subject. Once the ridge parameter was found, code with the MATLAB TRF toolbox was used to calculate mTRFs for each of the speech representations (code available at https://github.com/mickcrosse/mTRF-Toolbox). The TRFs were trained on 19 of the trials and used to predict the EEG responses to the remaining trial. Training and testing steps were repeated until all trials had been left out and used as the test trial.

### Assessment of Model Performance

The Pearson’s correlation coefficient, *r*, between predicted and measured EEGs was the dependent measure used to assess how well other feature vectors were represented in the EEG. As such, *r* was used to compare EEG prediction accuracy between the different TRF models. A baseline prediction accuracy was defined as the *r* value between the EEG predicted when using an envelope-derived TRF and the corresponding measured EEG response. Improvement from this baseline correlation value created a metric for success when using IC-response TRFs to predict the EEG. Prediction correlation values for spectrogram- and AB-envelope TRFs were also included for further comparison. A repeated measures ANOVA test was used to compare distributions of prediction correlation values across the different TRF models. Post-hoc comparisons between TRF models were done using Bonferroni corrected paired t-tests.

## Results

High-density (128 channels) EEG responses were collected from 19 subjects as they listened to excerpts from an audiobook (*The Old Man and the Sea* by Ernest Hemingway) containing narrative speech from an American male speaker. We used these responses and several features derived from the speech heard by the participants to fit TRF models and predict unseen EEG. In particular, we were interested to explore whether simulated IC responses driven by the stimulus could better predict the EEG compared to previously used features derived directly from the stimulus. Pearson’s correlation coefficients, *r*, between the predicted and measured EEGs were used to assess how well each TRF model predicted the EEG.

Twelve electrode channels over the frontocentral region of the scalp were used for analysis (Fig. 4, blue dots). Analysis channels were chosen based on examining the distribution of prediction correlations across the scalp for all four TRF models. The distributions were not significantly different (p>0.05, ANOVA) allowing for a single set of 12 electrodes with the highest prediction correlations to be chosen that did not bias the results towards any of the models (consistent with the approach in DiLiberto et al., 2015).

**Figure 4:**
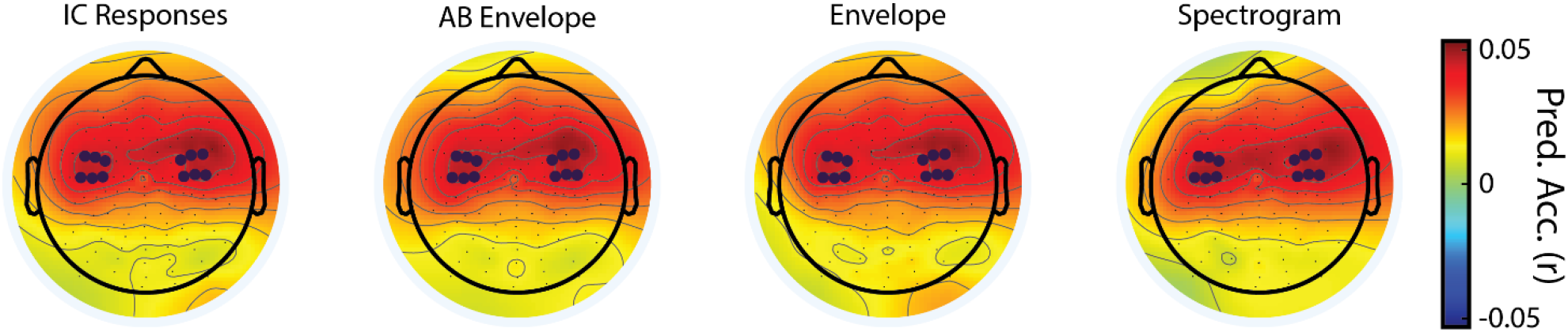
Topographical representation of prediction correlations between predicted and actual EEGs. Topographical distribution of prediction correlation values plotted onto a schematic diagram of the scalp. The electrodes used in analysis are emphasized in dark blue. The colormap indicates the Pearson correlation between the predicted and recorded (unaveraged) EEG.

### Comparing Performance of Feature-Specific TRFs

The grand means of prediction correlation values were compared across the different TRF models (Fig. 5A). We first performed a one-way repeated-measures ANOVA with factor TRF model which revealed a significant main effect (F_(4,72)_=30.87, p=5.8×10^−15^), indicating that some models were better at predicting EEG than others. The IC-response mTRF outperformed the envelope and spectrogram mTRFs (IC vs Envelope: T_(18)_ = 4.63, p = 3.6×10^−7^; IC vs.

**Figure 5:**
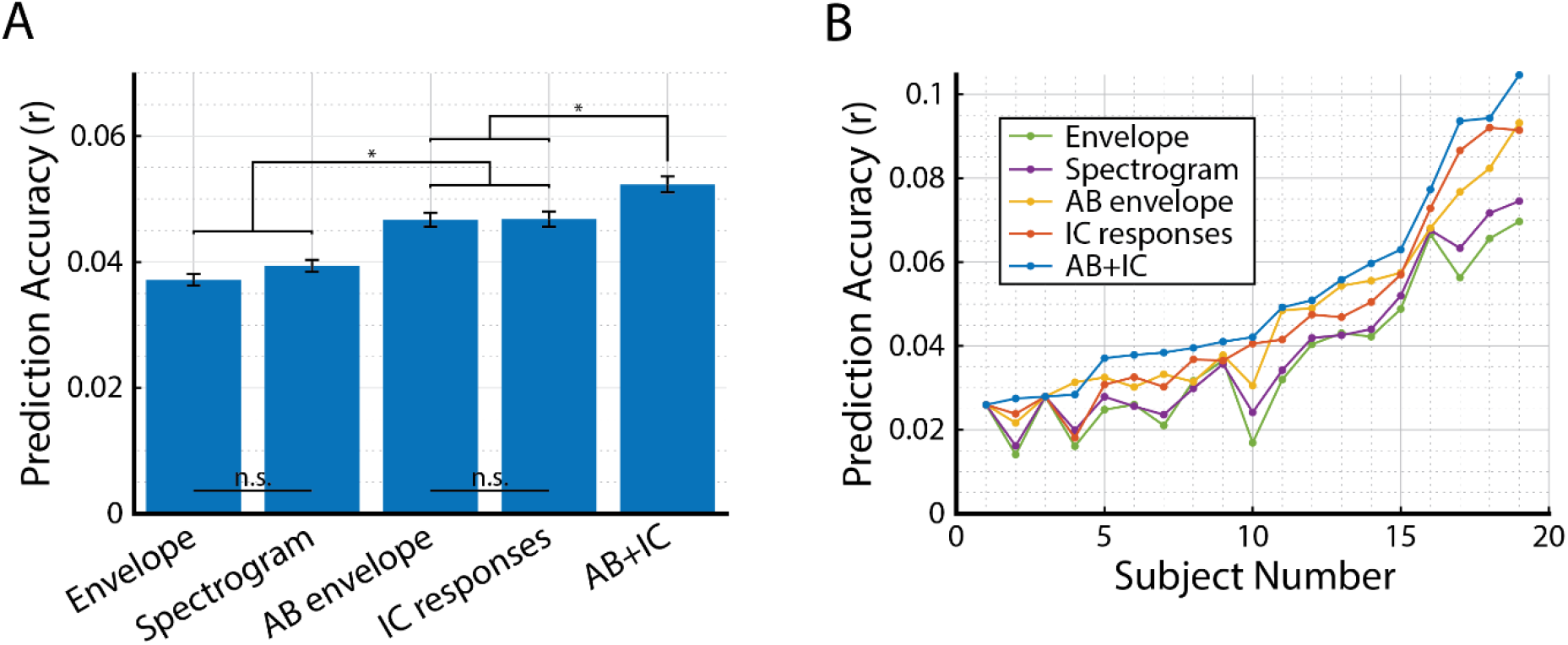
Comparison of IC-model derived mTRF to acoustic feature derived TRFs. **A)** The grand mean prediction correlation values for each type of TRF (mean ±SEM). The IC-model and AB-envelope TRF models were significantly higher than the envelope and spectrogram models. There was no significant difference between envelope and spectrogram prediction accuracy or between IC-model and AB-envelope prediction accuracy (p>0.05). The combined AB+IC TRF outperformed all other models. **B)** Correlation values were plotted for each subject individually. Data was sorted according to the prediction correlation values for the AB+IC model. Despite some variability across subjects, it is clear that the AB+IC mTRF model performs the best and the envelope TRF has the poorest performance across subjects.

Spectrogram: T_(18)_ = 4.63, p = 4.5×10^−7^). The IC-response mTRF performance was not significantly different from the AB-envelope-derived TRF (T_(18)_ = 0.062, p>0.5).

Because both the AB envelope and IC response predicted the EEG with comparable performance, we explored whether the two speech representations predicted different aspects of the EEG signal. To test this hypothesis, we analyzed the correlation between the predictions derived from each of these models and compared that to the correlation between envelope- and spectrogram-based prediction. For comparison, the EEG predicted from the envelope-derived and spectrogram-derived TRFs were strongly correlated (r=0.79), suggesting that these TRFs predict similar features of the EEG signal. In contrast, there was a significantly weaker correlation between the AB-envelope and IC predicted EEGs (r = 0.59; T_(18)_ = 6.59, p = 3.4×10^−6^), suggesting that these two speech representations were capturing more complementary aspects of the EEG compared to envelope and spectrogram model. Given this evidence, we fit a joint model (AB+IC) and hypothesized that it would predict both aspects of EEG, thus improving predictions of the overall EEG signal. As expected, the combined AB+IC mTRF predicted the EEG significantly better than its constituent features (Fig. 5A; AB+IC vs AB: T_(18)_ = 4.82, p = 0.0064; AB+IC vs. IC: T_(18)_ = 6.77, p = 0.0076).

### Analyzing Inter-subject Variability

To better assess model performance given inter-subject variability, the recorded *r*-values for each TRF-model were plotted individually and compared across subjects (Fig. 5B). Although the results show variability across subjects, the AB+IC mTRF model produced higher correlation coefficients than all other TRF models across 18 of 19 subjects. Further, the envelope- and spectrogram-TRF models consistently performed the poorest of the five models.

## Discussion

Auditory stimuli such as speech undergo substantial processing as they ascend from the cochlea to the cortex. Despite awareness of such transformations and despite the ability to extract subcortical responses to continuous speech (Forte et al., 2017; Maddox and Lee, 2018; Polonenko and Maddox, 2021) to our knowledge subcortical processing above the level of cochlear filters (Kulasingham et al., 2020; Gillis et al., 2021; Weineck et al., 2022) has not been incorporated into EEG analyses of continuous speech. Here, we have shown that including subcortical processing, in the form of auditory-midbrain model responses, allows for the derivation of TRFs that predict EEG responses with higher accuracy than previous TRFs based on acoustic features derived directly from the speech stimuli.

More specifically, in this work, the SFIE midbrain model was used to produce model IC responses to speech which, in turn, were used to derive TRFs for predicting EEG responses. The Pearson’s correlation coefficient between predicted and measured EEG was used to evaluate if incorporating the IC responses into the mTRF pipeline could produce better EEG predictions than those based on the envelope, spectrogram, or AB-envelope derived from the speech. Such acoustic features have previously been reported as successful methods for predicting EEG (Lalor et al., 2009, Ding and Simon, 2012, DiLiberto et al., 2015; Drennan and Lalor, 2019). However, those approaches generally ignore the substantial amount of processing that occurs before the input signal reaches the cortex and influences the EEG signal. Thus, incorporating a model of the IC into the pipeline could lead to better predictions of the EEG. Similar hierarchical, predictive models have been implemented, for example, in the nonhuman primate visual neuroscience literature (Mineault et al., 2012). As expected, the IC-response TRF outperformed the envelope and spectrogram TRF models; but, it did not predict the EEG better than the AB envelope-derived TRFs. However, the IC-response and AB-envelope-based predictions were not as highly correlated with each other as envelope and spectrogram predictions and combining IC responses and the AB envelope features into one mTRF model provided the best of the tested mappings to EEG responses. This suggests that the IC responses are capturing additional information beyond that which was obtained directly from the stimulus and supports our original hypothesis.

IC neurons lend themselves well to TRF analyses of speech encoding, as most cells are rate-tuned to both audio frequency and amplitude-modulation (AM) frequency. The display of spectral tuning is often characterized by a strong sensitivity to a certain frequency, or best frequency (BF), while low-frequency AM tuning is often characterized by a best modulation frequency (BMF; Krishna and Semple, 2000; Joris et al., 2004; Nelson and Carney, 2007). Further, a majority of IC BMFs fall within the range of voice pitch (Langner, 1992) making them suitable for analyzing speech stimuli (Delgutte et al., 1998; Carney et al., 2015). BMFs are best represented by the peaks (or troughs) in MTFs, which depict average discharge rate as a function of AM frequency and can be classified as either band-enhanced, exhibiting an increased discharged rate at BMF, or band-suppressed, exhibiting a decreased firing rate at BMF. The SFIE model simulates responses from both cell types, providing a robust population response that is representative of the midbrain encoding of complex sounds. Given this evidence, it is not surprising that the IC-response mTRF was able to represent the EEG significantly better than previous models, which typically only incorporate acoustic features of the speech stimuli (albeit sometimes passed through a very simple gammatone filter model of the cochlea). In particular, it is likely that the addition of AM tuning properties, typically present at the level of the midbrain provides, has provided much of the improvement in modeling the EEG.

The analysis presented in the current work focused specifically on EEG responses to speech. This was a natural choice given the importance of speech in everyday life, as well as the wealth of previous research aimed at modeling EEG responses to natural, continuous speech (Lalor and Foxe, 2010; Ding and Simon, 2014; Myers et al., 2019; Brodbeck and Simon, 2020). And while the inclusion of the IC-model within the framework has improved our EEG modeling, it may be that the use of speech in particular may actually have produced more modest benefits than would have derived for other types of audio stimuli. This is because envelope/spectrogram modeling has already been shown to work quite well for speech in particular – given the large amplitude modulation depths seen in natural speech and the importance of envelopes for speech intelligibility in general (Shannon et al., 1995; Smith et al., 2002). Moreover, the frequency content of speech – while obviously very important – does not tend to vary all that much for individual speakers. In particular, amplitude modulations across frequencies will tend to be highly correlated, limiting the impact of adding the ICresponses with its ability to capture AM information. As such, it might be the case that the framework we have introduced here – including an IC model in an EEG modeling pipeline – would produce larger benefits in the context of other stimuli. In particular, greater improvements in EEG prediction accuracy might derive for signals with a more heterogeneous pattern of amplitude modulations across frequencies, like music. Indeed, EEG tracking of the envelope of music has often been shown to be much weaker than for speech (Zuk et al., 2021). Future work will apply the framework presented here to modeling EEG responses to music.

Another interesting possible use of the framework presented here could be for the refinement of auditory subcortical models themselves. Although models of the auditory periphery have been used successfully to predict human speech perception (Heinz, 2010; Moncada-Torres et al., 2017; Bruce, 2017; Zaar and Carney, 2022), such models are often fit using data recorded from non-human mammals (Carney, 1993; Zhang et al., 2001; Zilany and Bruce, 2006, 2007; Zilany et al., 2009, 2014). While these models should work well given evolutionary homologies in the midbrain (Webster, 1992; Grothe et al., 2004; Woolley and Portfors, 2013), it is also true that speech is a particularly special signal for humans. As such, the processing of speech by humans involves predictions (Kutas and Hillyard, 1980, 1984; Leonard et al., 2016; Zoefel, 2018; Broderick et al., 2018) and attention (Cherry, 1953; McDermott, 2009; Mesgarani and Chang, 2012; Golumbic et al., 2013; O’Sullivan et al., 2015) effects that are unlikely to be present in non-human animals. One could imagine constraining and refining parameters of a subcortical model based on EEG prediction accuracy to have those subcortical models better capture human-specific subcortical auditory processing. Of course, validation of such models would be extremely important given the relatively low SNR of EEG and the risk of overfitting, and could perhaps be carried out using intracranial recordings in human neurosurgical patients.

In sum, we have shown that incorporating a well-established model of IC neuronal activity can improve models of EEG responses to natural speech. Given the relatively low SNR of EEG, any improvements in the ability to model that EEG could have important benefits in electrophysiological research on speech and language processing. Future work will aim to extend the framework to other auditory stimuli.

## Acknowledgements

This work was supported by NIH-DC010813 and the Del Monte Institute for Neuroscience at the University of Rochester.

